# Stereo In-Line Holographic Digital Microscope

**DOI:** 10.1101/790535

**Authors:** Thomas Zimmerman, Nick Antipa, Daniel Elnatan, Alessio Murru, Sujoy Biswas, Vito Pastore, Mayara Bonani, Laura Waller, Jennifer Fung, Gianni Fenu, Simone Bianco

## Abstract

Biologists use optical microscopes to study plankton in the lab, but their size, complexity and cost makes widespread deployment of microscopes in lakes and oceans challenging. Monitoring the morphology, behavior and distribution of plankton *in situ* is essential as they are excellent indicators of marine environment health and provide a majority of Earth’s oxygen and carbon sequestration. Direct in-line holographic microscopy (DIHM) eliminates many of these obstacles, but image reconstruction is computationally intensive and produces monochromatic images. By using one laser and one white LED, it is possible to obtain the 3D location plankton by triangulation, limiting holographic reconstruction to only the voxels occupied by the plankton, reducing computation by several orders of magnitude. The color information from the white LED assists in the classification of plankton, as phytoplankton contains green-colored chlorophyll. The reconstructed plankton images are rendered in a 3D interactive environment, viewable from a browser, providing the user the experience of observing plankton from inside a drop of water.

## 1. INTRODUCTION

Plankton are a large class of aquatic microorganism which are unable to swim against a current and vary significantly in size and features. Plankton make up the bottom of the aquatic food chain and produce up to two-thirds of atmospheric oxygen [1] and about half of the primary food production [2]. They are also the largest sequester of carbon, removing carbon dioxide from the air through photosynthesis and storing it at the bottom of the ocean [3]. These vital roles make maintaining the health of plankton essential. Recognizing their vital role, scientists have been collecting and monitoring plankton in the environment. Traditionally this has been done with nets or bottles to capture the small creatures and examine them in a laboratory under a microscope [4]. Advancements in image sensors and electronics have made it possible to place microscopes *in situ*, to investigate the relationship between plankton and its environment. These microscopes are custom built in small volume from specialized parts making them relatively expensive, limiting the number that can practically be deployed in the field. This is a severe limitation, as plankton is distributed all over the globe, which makes it difficult to estimate species richness and diversity [5] and generalize from local to global relationships.

The goal our research is to leverage commercial off-the-shelf (COTS) electronics and components to reduce the cost of *in situ* plankton microscopes to capture and analyze plankton shape, populations features and behavior. By making *in situ* microscopes inexpensive, more of them can be deployed, providing greater spatial coverage to gather a better understanding of plankton dynamic in a variety of conditions. To further reduce the cost and complexity of the microscope we implement lensless imaging [6] with a white light source to provide color information and direct in-line holography [7] to detect 3D structure and improve the resolution. Using COTS camera components presents several technical and system design challenges.

Direct in-line holographic microscopy (DIHM), first described by Gabor in 1947, eliminates the need for lenses, while offering high resolution images, down to the scale of bacteria [8]. But to achieve such high spatial resolution, the laser is made highly coherent with an optical bandpass filter [9], pin hole or fiber optic [10], all of which increases the cost and complexity of the microscope. An inexpensive red laser has a sufficiently small aperture to create a holograph of plankton as shown in Figure 1.

**Figure 1.**
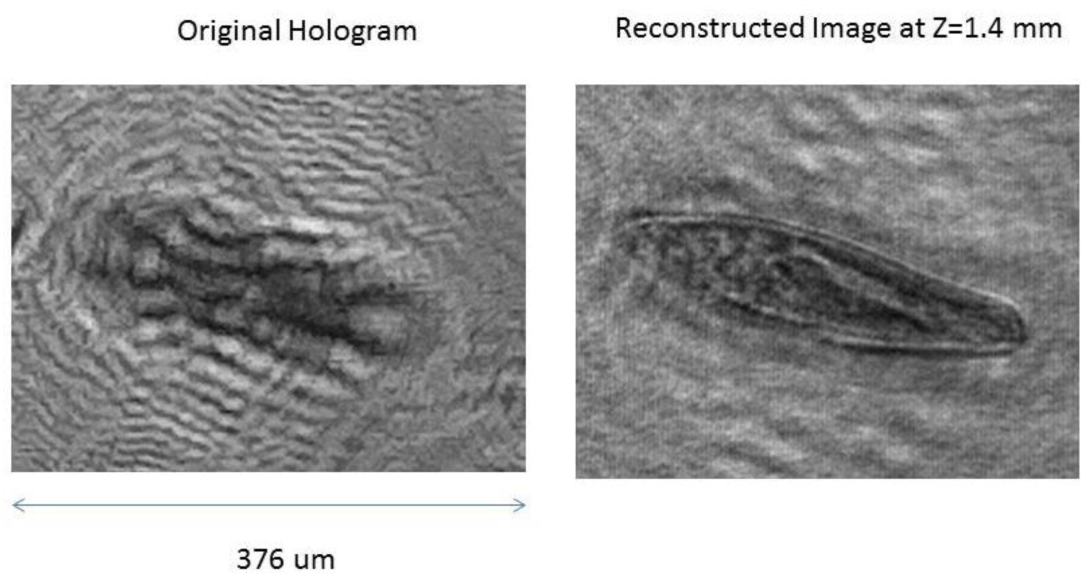
Hologram and reconstructed image of a paramecium taken with a consumer-grade red laser pointer (650 nm) captured with an OmniVision OV5647 image sensor (1.4 um pixel pitch). The laser is 25 mm above the image sensor. The plankton is swimming in ∼ 1mm deep drop of water on the image sensor’s protective glass cover.

Consumer digital cameras are designed primarily to take color picture of people and landscapes with little or no adjustments. The image sensors use rolling shutters which makes it difficult to dynamically change light sources at frame rate. In compact cameras, like those used in mobile phones, the image sensor and lens are molded together in plastic and are not designed for removal or modification. The image sensors are fabricated with a color filter on top of the silicon die which is extremely difficult to remove. In spite of all these challenges, the clear advantage of using an image sensor from a consumer camera is cost. For example, a modest 2 megapixel monochromatic image sensor appropriate for microscopy costs ∼$500 (Sentech). In contrast, the 5 megapixel color image sensor we are using costs about $7. Similarly, the monochromatic image sensor typically connects to a PC over USB3, often with proprietary software. In our system, the image sensor connects to a $10 ARM-based single board computer with a free Linux-based operating system that supports Python-based open-source modules.

In this paper we describe how we use COTS components to build a plankton microscope for studying the shape and behavior of plankton *in situ*. Observing plankton movement is important because disturbances in normal plankton predatory-prey behavior have been implicated as a factor in the production of harmful algal blooms [11]. Capturing plankton swimming has been achieved with video microscopes [12] and multiple exposures with a holographic microscope [13]. Many of the *in situ* microscopes are designed to be submerged, requiring a high pressure housing which greatly increases the cost. Our approach is to locate the microscope on or above the water surface and pump samples to the microscope, allowing the microscope housing to be constructed from inexpensive PCV pipes and caps.

## 2. MATERIALS AND METHODS

The plankton microscope includes the following system components; light source, image sensor, image capture and processing hardware, sample delivery and data presentation. Many of these components use high-volume commodity parts sourced directly from China (www.aliexpress.com) making them extremely inexpensive, as demonstrated by the bill of materials (Table 1). Each system component is discussed in the following sections.

**Table 1.** Bill of materials for stereo in-line holographic digital microscope

| Item | Cost |
| --- | --- |
| Image Sensor (Raspberry Pi Camera Version 1) | \$7.00 |
| Red Laser | \$0.30 |
| White LED | \$0.10 |
| Fiber Optic | \$0.10 |
| Raspberry Pi Zero W | \$10.00 |
| Peristaltic Pump | \$12.00 |
| Analog Comparator (LM393) | \$0.50 |
| Arduino Pro Mini Microcontroller | \$2.00 |
| **TOTAL** | **\$32.00** |

### 2.1 Light Source

Holographic microscopy creates high resolution images, providing shape features essential for plankton classification [14]. White light based lensless imaging provides color features, albeit at lower resolution, which is useful in distinguishing phytoplankton (plant-based) from zooplankton (animal-based) [15]. A 2D hologram contains 3D spatial information which is reconstructed at discrete distances from the image sensor (z). Our innovation is to use two light sources to triangulate the location of objects in 3D [16], limiting the reconstruction calculations to a subset of the imaged volume, and use a white LED and red laser as the light sources, producing a lower resolution color image and higher resolution monochromatic image, respectively.

The two light sources are located 25 mm above the image sensor, mounted in a black PVC cap. The light sources are 10 mm apart to create the disparity required for triangulation. The collimating lens of the laser is removed to reduce the intensity, which would otherwise saturate the image sensor. This has the added benefit of producing a wide beam, simplifying the alignment of the laser with the image sensor. A 0.3 mm diameter plastic fiber optic couples the white LED to the cap (Figure 2A), approximating a point light source. A 10-turn potentiometer in series with each light source is used to balance the brightness of each light source.

**Figure 2.**
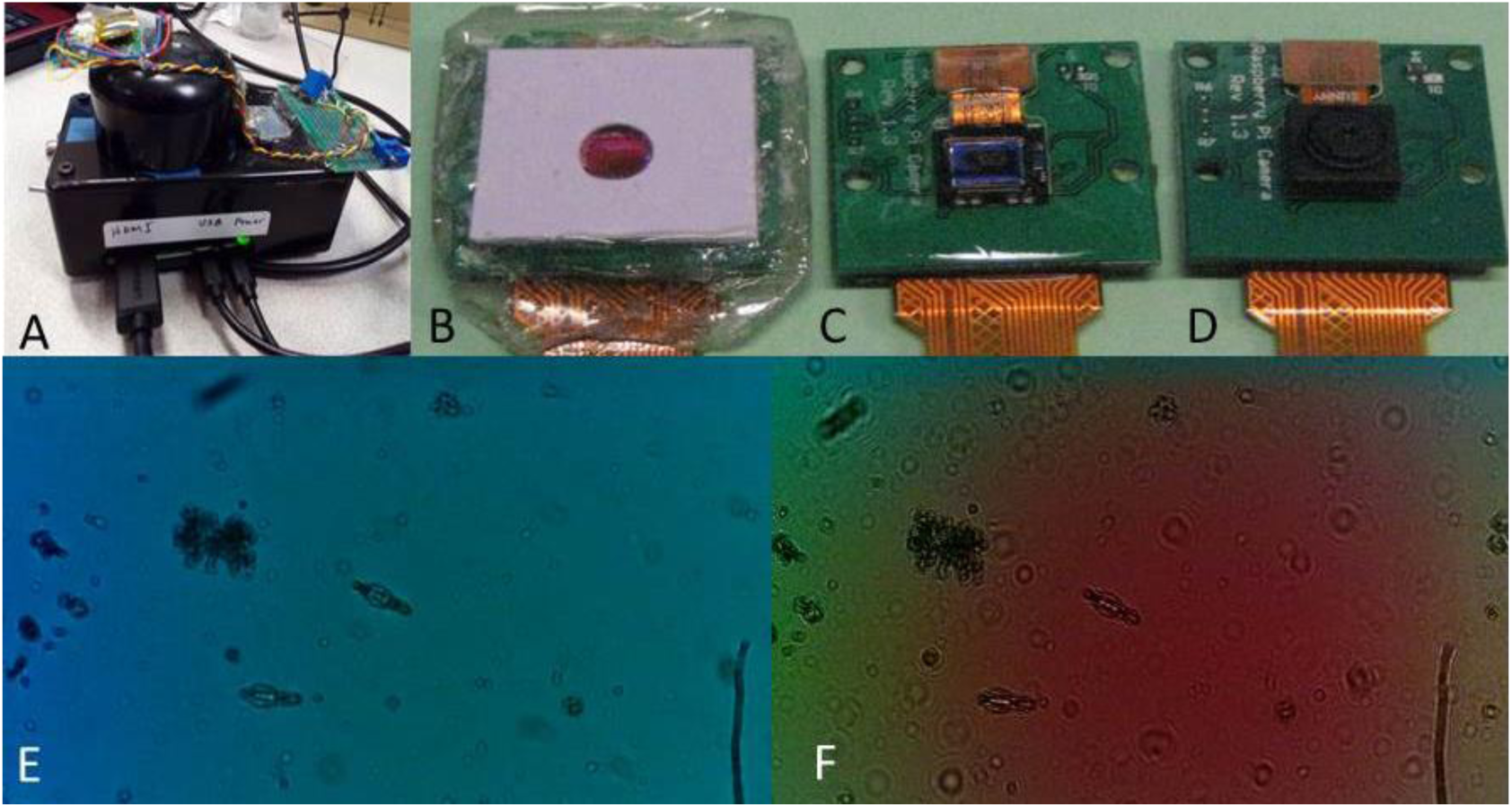
Prototype stereo microscope (A) contains a white LED coupled to a 0.3 mm diameter plastic fiber optic and a red laser light source 10 mm apart and 25 mm above an image sensor protected by a plastic cover (B). The image sensor (C) is prepared by removing the plastic lens and housing of a digital camera (D). The light sources are sequentially energized, synchronized to the frame rate to create a pairs of disparity stereo images, from the white LED (E) and red laser (F).

### 2.2 Image Sensor

The image sensor is extracted from a Raspberry Pi camera Version 1 (Figure 2D) which contains a 5 megapixel color image sensor (OmniVision OV5647) capable of streaming HD video (1920×1080) @ 30Hz. The plastic lens and the upper part of the plastic case is removed using diagonal cutters, exposing a 6.5 x 6.5 mm glass cover (Figure 2C) that protect the delicate gold bonding wires, which establishes the minimum distance a plankton sample can be from the image sensor.

The imager’s glass cover is not hermetic, so it must be sealed to prevent sample water from contacting the image sensor’s bonding wires. A 5.5 mm diameter hole, slightly larger than the imaging area (3.8 × 2.7 mm), is punched in a piece of plastic (PVC 0.8 mm thick) and bonded to the glass with silicone sealer (Figure 2B). Blue painter’s tape is temporarily placed on the glass above the imaging area during assembly and removed after the silicone sealer has set (24 hours), to preserve the optical clarity of the glass. The hole creates a well that is filled by the water sample to be imaged.

### 2.3 Image Capture Hardware

The prepared image sensor is connected by flex cable to the camera port of the Raspberry Pi Zero W board. Since no electrical modifications are made, the image sensor is compatible with all Pi camera software. Video is captured with the *raspivid* utility program which is preinstalled in the Raspbian operating system. The *picamera* library enables custom image capture code written in Python.

The Pi hardware and operating system provides user interface (e.g. mouse and keyboard input, HDMI output), WiFi connectivity, storage (USB flash and disk), and general purpose input/output (I/O) pins for controlling external hardware. However it is not responsive enough for controlling the frame-based switching of light sources, as reading and writing the I/O pins exhibits several milliseconds of jitter. To address this limitation, an inexpensive microcontroller (Arduino Pro Mini) is used to control the sequencing of the light sources.

The beginning of each video frame is detected by analyzing the clock signal of the Camera Serial Interface (CSI), accessible on the Raspberry Pi board. The clock signal consists of short pulses indicating row scans and a long pulse indicating the start of a new frame. One section of a dual analog comparator (LM393) level shifts the ∼1 volt clock signal to 5 volts and the other section implements an RC pulse-width circuit to detect the long pulse that indicates the beginning of a frame. This signal is received by the Arduino which alternately drives the red laser and the white LED for a fixed duration (e.g. 10 msec).

### 2.4 Image Processing

The image processing pipeline begins by capturing a short 1080p video (e.g. 10 seconds) of plankton, generating a sequence of frames alternately illumined by red laser and white LED. The illumination duration, image sensor exposure time and gain are set to accommodate the red laser and white LED brightness and to prevent light from one frame from exposing pixel rows of the next frame, due to the nature of a rolling shutter [17]. In our setup the combination of a light duration of 10 msec, exposure time of 39.2 msec and ISO (gain) of 800 satisfies these conditions.

The captured video is separated into pairs of 1920×1080 images. For each image in the pair, plankton is detected by applying a threshold (determined empirically) to create a binary image. A bounding box is established around each plankter using the OpenCV library function *findContours*. Bounding boxes in holographic images are doubled in width and height to assure the capture of fringes. Each plankter in the white LED image is matched by correlation to its corresponding disparity image in the red laser image, determined by differencing their binary images. The correlation test is applied only to plankter within an expected search area to reduce computation. The distance between the centers of the disparity pair is proportional to the height of the plankter above the image sensor.

Each plankter must be tracked across frame boundaries to enable swimming behavior analysis. For each plankter in a frame, we examine the next frame and find the nearest plankter in three dimensions. This simple method performs well due to the high frame rate and sparsity of plankton occurring in nature. However, lab cultured samples have a much higher plankton density which introduces collisions that confuse the simple tracker. For these conditions we use a more sophisticated tracker that includes matching plankton shape and velocity.

### 2.5 Holographic Reconstruction

In the holographic imaging geometry of the microscope, plankton in the water are a negative distance, -*Z*, away from the sensor plane. Because the red laser is approximately spatially coherent (with center wavelength λ), the measured intensity at the sensor is well modeled using scalar wave propagation. Assuming the laser distance is an infinite distance away, the sensor-plane intensity is

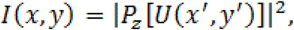

where *U*(*x′,y′*) is the complex sample transmittance and *P*_*Z*_ is coherent propagation by distance *Z*. The propagation operator is the well-known angular spectrum propagator:

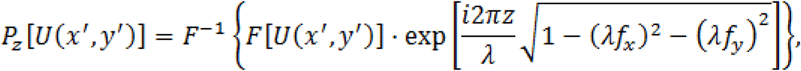

where *f*_*x*_ and *f*_*y*_ are the x- and y-direction spatial frequencies, and *F* is discrete Fourier transforms [18]. By convention, when 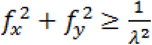, the complex exponential term is set to zero. Because the phase information of the electric field at the sensor is destroyed by the magnitude-squaring operation, it is not possible to directly compute *U*(*x′,y′*). However, it is possible approximate qualitatively the sample’s intensity, |*U*(*x′,y′*)|^2^, enough to improve the identifiability of the sample. By making the common inline holography approximation that the phase of the field at the sensor is zero, the field amplitude is approximately 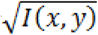. Though this is not physically true, it can be interpreted as assuming that the field at the sensor has been produced by two fields of identical amplitudes and conjugate phases, one a distance from the sensor and the other at –*Z* The sample’s intensity can then be approximated by propagating the measured amplitude a distance –*Z* and taking the magnitude-squared:

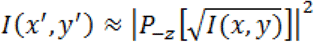

Due to the zero-phase assumption, a second image of the twin object at +z, propagated effectively by −2z, corrupts the object intensity estimate. However, the visual degradation due to the twin image is tolerable because it is quite blurry after propagating a long distance. The image sensor’s protective glass cover establishes the minimum distance between the sensor plane and the plankton, providing a sufficiently blurred secondary image. The image reconstruction function written in Python is provided in the Appendix.

### 2.6 Sample Delivery

Phytoplankton requires sunlight to perform photosynthesis so they are limited to the euphotic zone which can reach several hundred meters. Considering that water pressure increases one atmosphere per 10 meters, plankton microscopes typically are housed in metal vessels. To reduce cost, we rely on a tube submerged to the desired sampling depth to bring water up to the microscope, avoiding the need for an expensive high-pressure enclosure. A peristaltic pump inside the microscope draws up sample water which washes over and fills the sampling hole (Figure 2B). The pump is energized periodically (e.g. on for 10 seconds every 5 minutes) to wash off the previous sample and load a new sample. This must occur frequently enough to prevent the water from drying out, which would otherwise create permanent deposits (e.g. salt) on the imager’s glass cover. The resulting sample volume captured in the well is about 24 uL (5.5 mm diameter x ∼ 1 mm deep) while the actual viewing volume is about 10 uL (3.8 × 2.7 mm × ∼1 mm deep). This allows plankton to move freely in the larger sample volume so swimming behavior can be observed.

### 2.7 Data Presentation

We have created a web-based viewer (Figure 3) for the rendering an interactive 3D scene of the recorded video. The viewer runs on any web browser compatible with HTML5 and WebGL API (e.g. Firefox, Chrome and Edge). The rendering is managed through the library JavaScript 3D library *Three*.*js*, creating a virtual environment in which the user has full mobility to observe from different points of view, providing the visual experience of being inside the drop of water. To further increase the level of immersion, support for virtual reality interaction has been implemented. The application is natively based on a client-server model. The server manages requests from the client for cropped images of plankton and their 3D location, based on their contour and bounding box, respectively. The client renders the 3D scene by creating sprites from the cropped images and places them in the scene based on their 3D location and the user’s viewing perspective. The user dynamically controls their perspective by mouse and keyboard for a computer, touch gestures and tilt for a mobile phone, or head tracking for a head mounted display (e.g. Oculus Rift).

**Figure 3.**
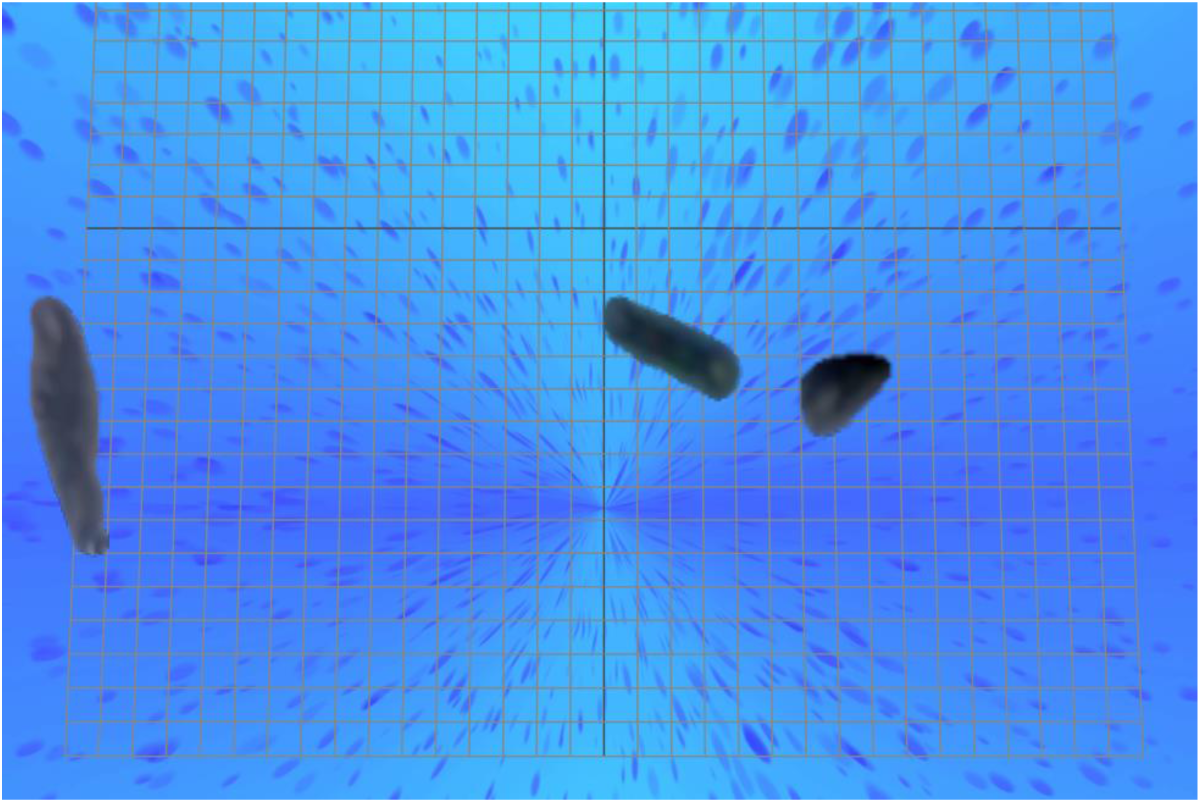
Screen capture of a browser-based viewer of real plankton placed in an interactive 3D virtual environment. Cropped images of plankton are rendered in the three dimensional space based on their location in the microscope well and the user’s viewing perspective controlled by input devices including mouse, keyboard, mobile phone or VR headset tracking.

## 3. RESULTS AND DISCUSSION

To determine the prediction accuracy of plankton height (z) above the image sensor, the predicted and actual z for 31 plankton samples are collected. A video of plankton is recorded and the resulting holograms for the 31 samples are reconstructed in 100 um steps (Figure 4). The reconstructions are manually reviewed and z corresponding to the best-looking reconstruction is selected. Another sequence of reconstructions is performed on the samples at 10 um steps around the selected z to establish the actual z value. The disparity is determined for each of the samples to produce the predicted z values. A regression analysis for the 31 samples indicates that z value prediction by the disparity has an R2 value of 0.96 (Figure 5). Using the z value predicted by disparity and the x, y location determined by the bounding box constrains the reconstruction volume to a fraction of the sampling volume, significantly reducing reconstruction calculations.

**Figure 4.**
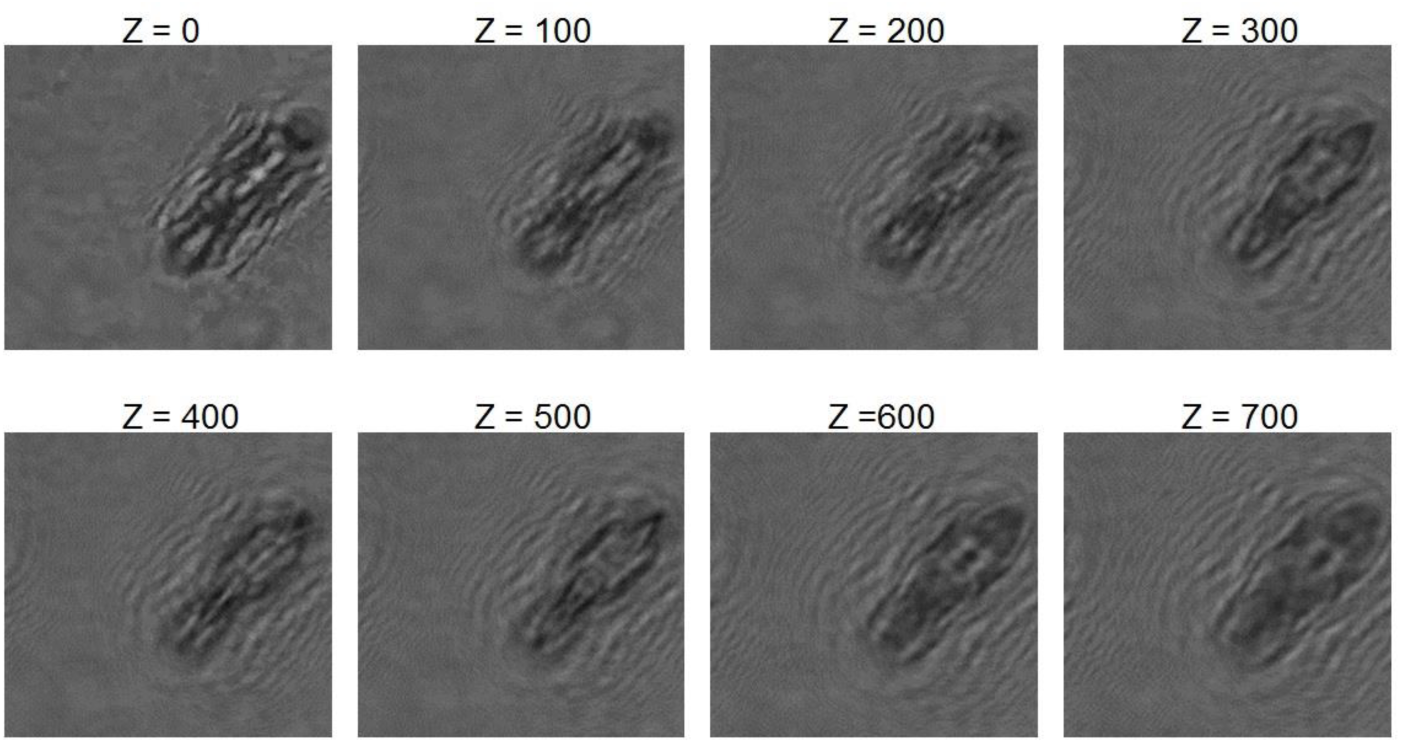
Holographic reconstruction of paramecium captured with a red laser at various heights above image sensor (Z).

**Figure 5.**
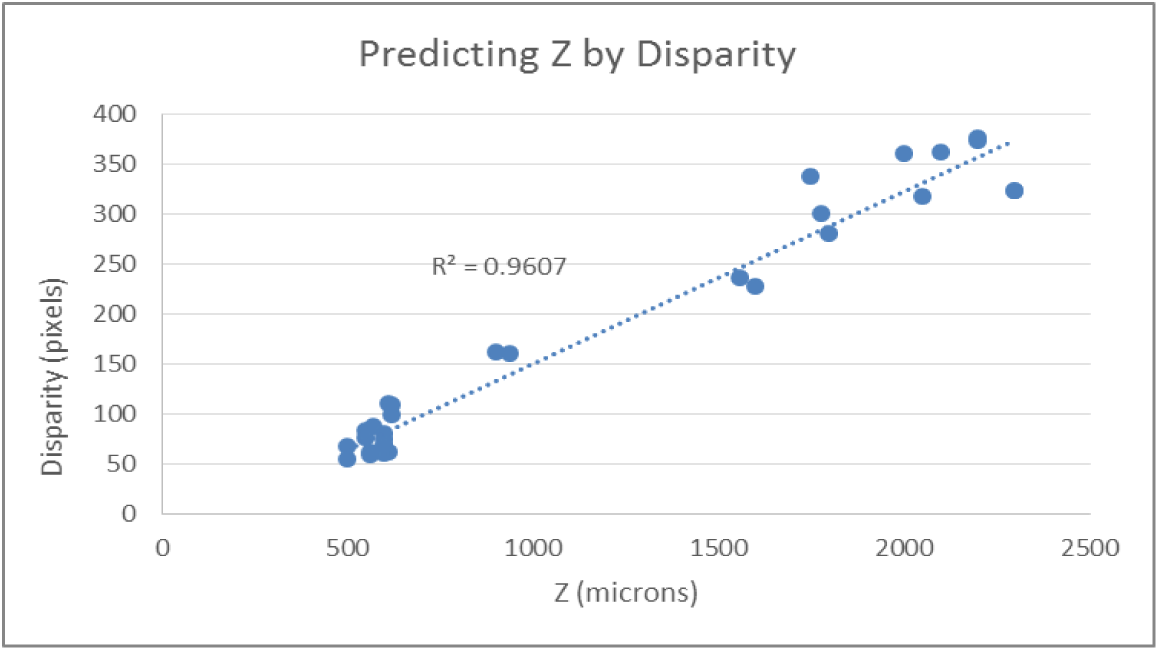
Height of plankton above the image sensor (z) is automatically predicted from the disparity of a pair of images produced by two sequentially illuminated light sources (Y axis). The measured distance (X axis) is determined by selecting the best holographic reconstruction.

In conclusion, a microscope that combines white-light lensless imaging and direct inline holography is implemented using inexpensive commodity components. The combination of the two imaging modalities provides low-resolution color and high-resolution spatial information for classifying and studying plankton behavior, while decreasing computation demands. The Raspberry Pi platform provides inexpensive hardware and an open-source community of code, documentation and support. The operating system contains a Python integrated development environment (IDE), providing not only a microscope, but a platform for further development and customization.

## ACKNOWLEDGMENTS

This material is based upon work supported by the National Science Foundation under Grant No. DBI-1548297. Disclaimer: Any opinions, findings and conclusions or recommendations expressed in this material are those of the authors and do not necessarily reflect the views of the National Science Foundation. Thomas Zimmerman, Sujoy Biswas, Daniel Elnatan, Jennifer Fung, Vito Paolo Pastore and Simone Bianco would like to thank the NSF for financial support. Alessio Murru gratefully acknowledges the Italian Ministry of University and Research for the financial support of his PhD scholarship (P.O.N. 2014-2020 Research and Innovation - FSE-FESR Ac. 1.1). We also would like to thank Joseph DeRisi of the University of California, San Francisco for suggesting the Raspberry Pi as the platform for the plankton microscope.

## Appendix

The following Python function receives a holograph image (im) and height (zdist) and creates a reconstructed image (imAmp).

def reconstructImage(im, zdist):

~~~
# im is monochromatic (one channel) holographic image, must have even number of rows and columns
# zdist is distance of object to image sensor (in microns)

input_img=np.sqrt(im)

# constants
dxy = 1.4e-6          # pixel spacing in micron
wvlen = 650.0e-9      # wavelength of light, 650 nm
# get image size, rows M, columns N, must be even values!
M, N = input_img.shape

# prepare grid in frequency space with origin at 0,0
_x1 = np.arange(0,N/2)
_x2 = np.arange(N/2,0,-1)
_y1 = np.arange(0,M/2)
_y2 = np.arange(M/2,0,-1)
_x = np.concatenate([_x1, _x2])
_y = np.concatenate([_y1, _y2])
x, y = np.meshgrid(_x, _y)
kx,ky = x / (dxy * N), y / (dxy * M)
kxy2 = (kx * kx) + (ky * ky)

# compute FT at z=0
E0 = np.fft.fft2(np.fft.ifftshift(input_img))   #ifftshift moves the center pixel to the upper left

# compute phase aberration
_ph_abbr = np.exp(−1j * np.pi * wvlen * zdist * kxy2)
output_img = np.fft.fftshift(np.fft.ifft2(E0 * _ph_abbr)) #fftshift moves the upper left pixel back to center.

# display reconstructed image
amp=np.abs(output_img)**2
imAmp=amp.astype(‘uint8’)
cv2.imshow(‘amp’,imAmp)

return
~~~

